# Recruitment of the local GABAergic population engages complex network interactions in the ventral subiculum

**DOI:** 10.64898/2026.10.06.757158

**Authors:** Mohsen Bahrami, Olivia Colarusso, Alyssa Danial, Jeff Weiner, Eva Bach

## Abstract

The ventral subiculum (vSub), a major output node of the hippocampal formation, has been implicated in diverse neurological disorders. Disruption of excitatory-inhibitory balance is a common feature of neurological disorders, with altered GABAergic signaling representing a critical component. More recently, the vSub has gained increasing attention as a component of neural circuits underlying emotional and motivational regulation, through its extensive connections with regions implicated in mental health disorders such as addiction and anxiety. Our previous work in models of alcohol use disorder and comorbid anxiety identified an important role for GABAergic inhibition in regulating vSub excitability, prompting us to investigate the functional consequences of recruiting the broader local GABAergic population. Although previous studies have examined individual interneuron subtypes, the network consequences of activating the global population of local GABAergic neurons remain poorly understood. To address this question, we expressed channelrhodopsin using an mDlx-dependent strategy to target GABAergic neurons in the ventral hippocampus and performed in vivo electrophysiological recordings from vSub neurons in anesthetized rats. Repeated activation of this GABAergic population at 20 Hz, a stimulation frequency commonly used in behavioral optogenetic studies, produced robust suppression of pyramidal neuron firing during stimulation. This direct inhibition was accompanied by network-dependent increases in post-stimulus activity that varied as a function of baseline activity, sex, and trial number. Complementary ex vivo electrophysiological experiments further supported network-dependent effects and revealed short-term synaptic plasticity that evolved across repeated stimulation trials, reflecting contributions from both pre-and postsynaptic mechanisms. Together, these findings demonstrate that recruitment of the broader GABAergic population produces effects extending beyond direct inhibition and can engage dynamic network responses that depend on stimulation history and circuit state. These findings highlight the importance of stimulation frequency, timing, and network context when interpreting the functional consequences of GABAergic circuit manipulation.

## Introduction

The subiculum is the major output structure of the hippocampal formation (O’Mara et al., 2001), and although it has traditionally been studied for its role in learning and memory, the ventral subiculum (vSub) in particular has increasingly been recognized as a critical substrate in the pathophysiology of a broad range of neurological and neuropsychiatric disorders. Clinical and preclinical evidence implicates vSub dysfunction in epilepsy, where it contributes to seizure initiation and propagation (Fei et al., 2021), as well as in a range of neuropsychiatric conditions, including schizophrenia, anxiety/stressor-related disorders, and addiction (Baset and Huang, 2024; Grace and Uliana, 2026; Herman and Mueller, 2006; Miller and Aoto, 2026). The vSub sends dense projections to a range of downstream targets implicated in emotional and motivational processes, such as the basolateral amygdala, nucleus accumbens, hypothalamus, and prefrontal cortex (Grace and Uliana, 2026; Wee and MacAskill, 2020). Because the vSub serves as a shared node linking hippocampal output to each of these circuits, even modest shifts in its local excitatory/inhibitory balance have the potential to be amplified into disproportionate effects on behavioral outputs and the corresponding pathologies.

Over the last several years, our lab has used rodent models to further characterize the vSub as a region of experience-dependent hyperexcitability relevant for understanding the pathophysiology of alcohol use disorder (AUD) and stress/anxiety-related negative emotional states that may exacerbate AUD symptoms. This line of inquiry has been motivated by clinical data which has demonstrated that patients with AUD have reduced hippocampal volume (De Bellis et al., 2000), specifically in the anterior hippocampus (Sullivan et al., 1995), the human brain region analogous to the rodent ventral hippocampus, compared to healthy controls. Likewise, anterior hippocampus activity has been associated with contextual, state-like anxiety in humans (Satpute et al., 2012). These clinical findings further support the epidemiological data highlighting the high comorbidity between AUD and disorders of negative emotion (Gilpin and Weiner, 2017; Oliveira et al., 2018). As such, we have used *ex vivo* whole-cell patch electrophysiology to identify and characterize the ventral hippocampus (and particularly the vSub) as a site of marked long-term synaptic adaptations as a consequence of chronic intermittent ethanol vapor exposure, a preclinical model of alcohol dependence which can also lead to anxiety-like behaviors (Bach et al., 2021b, 2021a; Bach and Weiner, 2025; Ewin et al., 2019). Notably, one of these studies revealed that the strength of local vSub GABAergic inhibition emerged as a key determinant of whether this vulnerability is expressed, with pharmacological blockade of GABA_A_ receptors sufficient to unmask an underlying hyperexcitable phenotype (Bach and Weiner, 2025).

Indeed, previous preclinical research has positioned GABAergic interneurons within the ventral hippocampus to actively shape the excitability of pyramidal neurons (for extensive review on interneurons in the hippocampal complex see (Pelkey et al., 2017)). Parvalbumin (PV) interneurons, for example, are recruited as animals transition into anxiogenic compartments of the elevated plus maze, and this recruitment coincides with a state-dependent reduction in PV-mediated inhibition of pyramidal neurons (Volitaki et al., 2024). Critically, inhibitory control is not monolithic, such that distinct interneuron populations appear to gate distinct pyramidal neuron ensembles to support different negative affective states (K. Li et al., 2024). Furthermore, naturally occurring individual differences in the strength of interneuron-pyramidal neuron coupling align with individual anxiety-related phenotypes, with low-anxiety animals exhibiting stronger inhibitory engagement of anxiety-related pyramidal neurons than their high-anxiety counterparts (Forro et al., 2022). Together, these findings establish that hippocampal interneurons are not simply a passive counterbalance to excitation, but a dynamically and specifically modulated feature of local circuit function with direct, causal consequences for behavior.

Although the vSub is a well-established site of plasticity in which local GABAergic interneurons dynamically shape pyramidal neuron excitability, investigations to date have focused on the specific and independent role of particular interneuron subtypes (Bratsch-Prince et al., 2026; Forro and Klausberger, 2023; Volitaki et al., 2024).

Because distinct interneuron populations can exert opposing or non-overlapping influences on pyramidal neuron activity (Lodge et al., 2023; Royer et al., 2012), manipulations restricted to a single subtype may not capture the net effect of local inhibitory tone on circuit output. As such, the functional consequences of recruiting the broader GABAergic population within the vSub remain uncharacterized both *in vivo* and *ex vivo*. To fill this gap, the current study leveraged optogenetic activation of the broader GABAergic population using an mDlx viral approach (Dimidschstein et al., 2016) and stimulation parameters commonly employed in behavioral optogenetic studies of awake, behaving animals. This approach allowed us to investigate the consequences of broad GABAergic recruitment at the level of individual neurons, local network organization, and synaptic mechanisms. Specifically, *in vivo* electrophysiology was used to characterize how repeated, direct recruitment of this population shapes the acute and longer-lasting firing properties of pyramidal neurons and to determine whether it disrupts the organization of ongoing network oscillations, while *ex vivo* electrophysiology was used to probe the underlying synaptic mechanisms.

## Methods

### Animals

Male and female Long Evans rats were purchased from Inotiv, IN and arrived between 6 and 10 weeks of age. Upon arrival rats were singly housed in clear cages (25.4 cm × 45.7 cm) and maintained on a 12:12 h light dark cycle. Rats had ad libitum access to food (Prolab RMH 3000, LabDiet: PMI Nutrition International, St. Louis, MO) and water throughout the study. Animal care procedures were carried out in accordance with the NIH Guide for the Care and Use of Laboratory Animals and were approved by the Wake Forest University Animal Care and Use Committee. All animal care and use procedures were carried out in compliance with ARRIVE guidelines.

### Surgical procedures

Following acclimatization male and female rats, weighing between 300-350 g and 200-230 g respectively, were anesthetized using sodium pentobarbital (40– 50 mg/kg, i.p.). For analgesic purposes animals were injected subcutaneously with 2 mg/kg meloxicam. The scalp was shaved and surgically scrubbed. Rats were placed in a stereotaxic frame and an incision was made centrally along the scalp. A craniotomy positioned directly above the posterior ventral subiculum was made bilaterally to allow microinjection needles to be lowered into the ventral hippocampus (vHC) targeting the ventral subiculum (stereotaxic coordinates in mm with male/female adjustments: AP: 5.85-6.2; ML: 4.5-4.75; DV: 6.4-7.1). Microinjection needles were used to transfect the vHC with 0.8–0.9 μL of a virus construct (pAAV9-mDlx-ChR2-mCherry-Fishell-3) expressing the excitatory opsin, channelrhodopsin (ChR2) at a rate of 0.1 μL/min (Addgene, Cambridge, MA). Microinjection needles were maintained in place for 5 min before being withdrawn. Animals used for in vivo and ex vivo electrophysiology were sutured and allowed to recover in their home cages.

### In vivo electrophysiology recordings

On the day of recording, animals were anesthetized with an initial loading dose of sodium pentobarbital (60 mg/kg for male rats and 50 mg/kg for female rats, i.p.).

Anesthetic depth was monitored continuously throughout the experiment, and supplemental booster doses of sodium pentobarbital (10 mg/kg, i.p.) were administered as needed to maintain a stable plane of anesthesia. The scalp was shaved and surgically prepared, and rats were positioned in a stereotaxic frame. A midline incision was made along the scalp, and a stainless-steel screw was implanted anterior to bregma to serve as a ground electrode, which was connected to the ground of the multielectrode probe. Bilateral craniotomies were performed directly above the posterior ventral subiculum (vSub). Multielectrode probes (NeuroNexus OA32LP-32), coupled with a juxtaposed optical fiber, were gradually lowered into the vSub. Recording locations were guided by general stereotactic coordinates (coordinate range based on 0.75 mm surrounding viral injection sites) and confirmed by progressing through the hippocampal formation along the dorsoventral axis, using established electrophysiological signatures of adjacent hippocampal regions to identify the vSub. In a subset of rats, recordings were obtained from multiple sites spanning distinct stereotactic coordinates within the vSub.

Acute spontaneous and optogenetically evoked neuronal activity was recorded from vSub neurons using an Open Ephys acquisition system. Raw voltage signals were sampled at 30 kHz and stored for offline analysis. Spike sorting was performed using Kilosort4, followed by manual curation using SpikeInterface and custom Python scripts. Units were classified as single units based on established criteria, including a signal-to-noise ratio (SNR ≥ 4), the presence of a clear trough surrounding 0–2 ms in the autocorrelogram (indicative of minimal refractory period violations), waveforms exhibiting spatial decay along the probe axis, and a unimodal distribution of spike amplitudes. Optogenetic stimulation of GABAergic neurons (mDlx⁺ neurons) was controlled using a custom Arduino-based stimulus initiator interfaced with a Master-8 pulse generator, which produced hardware-timed TTL outputs. Stimulation timing signals were simultaneously recorded via the Open Ephys I/O board. Optical stimulation was delivered through an implanted optical fiber using an LED driver (Doric LED driver 450 nM). Blue light (450 nm) was delivered as square pulses (1 ms pulse width) at 20-Hz for 5 s, with trials separated by 25 s, for a minimum of 20 trials. This stimulation paradigm enabled both phototagging of GABAergic neurons and targeted manipulation of this neuronal population to assess its impact on spontaneous activity in surrounding vSub circuitry. In total, 116 neurons were recorded across 14 animals (7 females, 7 males) for the completion of in vivo electrophysiological analyses.

### Group Comparison statistics

Firing-rate data were analyzed using linear mixed-effects models that accounted for the nested structure of neurons within recordings and animals. Fixed-effects structures included baseline firing rate, optical-stimulation epoch (before, during, and after), sex, and ramping subtype, with all relevant interactions evaluated. Significance of fixed effects was assessed using F-tests with Satterthwaite-approximated degrees of freedom, and estimated marginal means were used to characterize group-level differences and interaction patterns. To more accurately model the temporal dependence across successive trials within each neuron, residuals were modeled using a first-order autoregressive [AR(1)] covariance structure.

More specifically, let *FR_ijk_* denote the firing rate for trial *k* of neuron *j* within recording *i*, *Period_ijk_* denote the firing rate period, modeled as a categorical variable with baseline as the reference level and two indicator variables corresponding to stimulation (*period*^(1)^*_ijk_*) and post-stimulation (*period*^(2)^*_ijk_*) periods, *ITI_ijk_* denote the intertrial interval between trials *k* − 1 and *k* for neuron *j* within recording *i*, and *Trial_ijk_* denote the trial number, where *Trial_ijk_* ∈ {1,2,…,*N_ij_*}, and *N_ij_* is the total number of trials for neuron *j* within recording *i*. Then, the model can be written as follows:

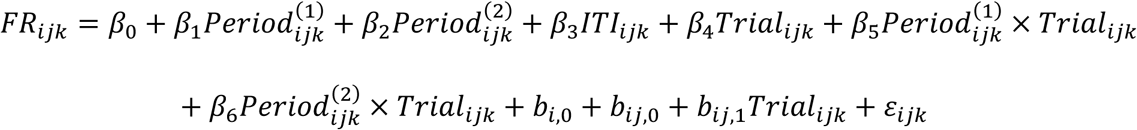

where *b_i_*_,0_ denotes the random intercept for recording *i*, *b_ij_*_,0_, *and b_ij_*_,1_ denote the neuron-specific random intercept and random slope for neuron *j* within recording *i*, *ε_ijk_* denotes the residuals. Random effects were assumed to have normal distributions:

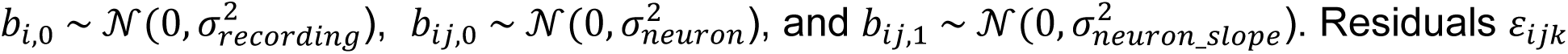

were modeled using a first-order autoregressive [AR(1)] structure within each neuron to account for temporal dependence across successive trials:

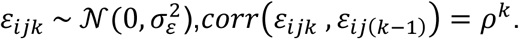

Multicollinearity diagnostics were assessed using standard diagnostics and no evidence of problematic multicollinearity was observed (see supplemental file for more details). All statistical modeling and post-hoc multicollinearity assessments were conducted in SAS (v 9.4) using the PROC GLIMMIX Procedure.

### GABAergic neuron classification

Neurons were classified as GABAergic based on their spike responses to optogenetic stimulation using a shuffle-based statistical framework designed to identify reliable, stimulus-locked firing while minimizing false positives arising from chance coincidences. For each stimulus presentation, spikes occurring within a 7-ms post-stimulus window were counted. Spike timing within this window was evaluated at 1-ms resolution, and a stimulus was considered responsive if at least one spike occurred within the window.

Stimuli were grouped into trials of 100 consecutive presentations. For each trial, the observed number of responsive stimuli was compared to a null distribution generated by randomly jittering stimulus times within ±20 ms for 1,000 iterations. This procedure preserved the overall spike train structure while disrupting precise stimulus– spike alignment. A trial-level p-value was computed as the proportion of shuffled iterations in which the number of responsive stimuli equaled or exceeded the observed value, and trials were considered significant at p < 0.01.

To classify neurons as GABAergic, statistical significance was combined with a reliability requirement to ensure consistent optogenetic responsiveness. Specifically, a neuron was classified as GABAergic only if at least 30% of trials were significant and, within each significant trial, at least 10% of stimuli elicited spikes. This dual criterion ensured that classification reflected reproducible stimulus-locked activation across trials rather than rare or isolated firing events, which is particularly important given the large number of stimulus presentations and the presence of neurons with low baseline firing rates.

### Quantification of mDlx-dependent inhibitory responses

Spike activity was quantified on a per-trial basis during three defined epochs: before stimulation, during stimulation, and after stimulation. For each neuron, firing rates (Hz) were calculated separately for each epoch across all trials. Mean firing rates and associated variability were computed across trials for each epoch.

To assess whether firing rates differed across epochs within a neuron, a Friedman test was used to perform a nonparametric repeated-measures comparison across the three conditions. When a significant main effect was observed (α = 0.05), post-hoc pairwise comparisons were conducted using Wilcoxon signed-rank tests between epoch pairs (before vs. during, during vs. after, and before vs. after). To control for multiple comparisons, a Bonferroni correction was applied, yielding a corrected significance threshold of α = 0.0167. All statistical tests were performed within neurons to evaluate stimulation-related changes in firing rate.

### Trial-Dependent Changes in Spontaneous Firing Following Stimulation

To assess changes in spontaneous firing across repeated trials, spike activity was analyzed during non-stimulated epochs surrounding each stimulus train.

Cumulative spike counts were computed for each trial to visualize trial-by-trial changes in spontaneous firing dynamics. For statistical analysis, spontaneous activity within each trial was quantified by binning spike counts and converting these values to firing rates, yielding an average trial-level measure of spontaneous firing frequency. To determine whether spontaneous firing changed systematically over the course of the experiment, trial-wise mean firing rates were tested for linear trends across trial number using linear regression. The slope of this regression provided a quantitative measure of the direction and magnitude of changes in spontaneous firing across trials.

For a subset of neurons, an additional prestimulus baseline (5-15 min in duration) recording was available in which no stimulation was delivered. This baseline activity was used to assess whether post-stimulus firing rates exceeded the neuron’s intrinsic spontaneous firing in the absence of stimulation. Baseline recordings were segmented into consecutive non-overlapping 5-s bins, and spike counts were converted to firing rates to generate a distribution of baseline spontaneous activity for each neuron.

Post-stimulus spontaneous activity was quantified using the same binning procedure applied to the post-stimulus periods (0–25 s following stimulus offset) across all trials as described above. To directly compare baseline and post-stimulus firing, firing rate values from baseline bins and post-stimulus bins were contrasted using a nonparametric permutation test on the difference in mean firing rates. The observed mean difference between post-stimulus and baseline firing rates was compared against a null distribution generated by randomly permuting bin labels across conditions.

### Analysis of Neuronal Phase distribution to Slow Oscillations

To assess the relationship between neuronal firing and ongoing network oscillations, local field potentials (LFPs) were analyzed at the unit-specific recording channel. LFP signals were resampled at 1 kHz and subsequently bandpass filtered between 0.4 and 2 Hz to isolate the slow oscillatory activity. The instantaneous slow-oscillation phase at each spike time was extracted using the Hilbert transform. Phase-locking strength was quantified using the mean resultant vector length (R), calculated as the absolute value of the circular mean of the spike phases: 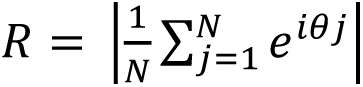 where represents the number of spikes and represents the slow-oscillation phase at the time of the spike. The statistical significance of phase locking was assessed using the Rayleigh test for non-uniformity of circular data for each epoch (baseline, stim and post-stim).

Additionally, differences in phase distributions across epochs were assessed using the Wheeler–Watson test for circular data in cells that had a minimum of 10 spike-specific phase observations within each epoch. Because phase observations contained tied values, pooled phase observations were assigned average ranks for tied observations, and the resulting ranks were transformed to circular angles for calculation of the Wheeler–Watson statistic. The test statistic was compared with a chi-square distribution to obtain the associated significance level.

### Ex vivo electrophysiology

Electrophysiological experiments were conducted following in vivo electrophysiological recording or from animals that were at least 6-weeks post mDlx viral injections. Animals were deeply anesthetized using isoflurane unless still experiencing a sufficiently deep anesthetic sodium pentobarbital mediated plane. Following decapitation, the brain was removed rapidly and suspended in ice-cold NMDG recovery solution containing in mM: 92 NMDG, 2.5 KCl, 1.25 NaH_2_PO_4_, 30 NaHCO_3_, 20 HEPES, 25 glucose, 2 thiourea, 5

Na-ascorbate, 3 Na-pyruvate, 0.5 CaCl_2_·2H_2_O, and 10 MgSO_4_·7H_2_O. NMDG was titrated to pH 7.4 with 17 mL ± 0.5 mL of 5 M hydrochloric acid(Ting et al., 2018). Transverse slices containing the vHC were cut at a thickness of 300 μm using a VT1000S Vibratome (Leica Microsystems). vHC slices were placed in a holding chamber containing NMDG recovery solution. Slices were allowed to recover for 35 min before being transferred to a chamber filled with artificial cerebral spinal fluid (aCSF) containing in mM: 125 NaCl, 1.25 NaH_2_PO_4_, 25 NaHCO_3_, 10 D-Glucose, 2.5 KCl, 1 MgCl_2_, and 2 CaCl_2_. NMDG recovery and aCSF holding solutions were oxygenated with 95% O_2_ and 5% CO_2_ and warmed to 32–34 °C. For recordings, a single brain slice was transferred to a chamber mounted on a fixed stage under an upright microscope (Scientifica SliceScope Pro 2000 microscope), where it was superfused continuously with warmed oxygenated aCSF. Whole-cell voltage-clamp recordings were made in presumptive pyramidal neurons of the ventral subiculum (vSub) using recording pipettes pulled from borosilicate glass (open tip resistance of 3–6 MΩ; King Precision Glass Co., Claremont, CA). The pipette solution contained (in mM): 130–140 Cs-gluconate, 10 HEPES, 1 NaCl, 1 CaCl_2_, 3 CsOH, 5 EGTA, 2 Mg^2+^-ATP, 0.3 GTP-Na_2_ and 2 Qx-314.

Intracellular Cs^+^ was used as the primary cation carrier in voltage-clamp recordings to block K^+^ currents, including postsynaptic GABA_B_ receptor-mediated currents, in the recorded neuron. Neurons in the ventral subiculum (vSub) were targeted for recording under a 40 × water-immersion objective (numerical aperture = 0.8) with infrared-differential interference contrast (IR-DIC) optics, as described previously (Bach et al., 2021). Electrophysiological signals were obtained using a Multiclamp 700B amplifier (Molecular Devices, Union City, CA), low-pass filtered at 3 kHz, digitized at 10 kHz, and recorded onto a computer (Digidata 1440A, Molecular Devices) using pClamp 10.0 software (Molecular Devices). Series resistance, measured from brief voltage steps applied through the recording pipette (5 mV, 5 ms), was < 25 MΩ and was monitored periodically during the recording. Recordings were discarded if series resistance changed by > 20% over the course of the experiment. Each recorded neuron represents an individual data point; recordings were made from a total of. To selectively stimulate terminals originating from the mDlx^+^ neurons synapsing onto vSub neurons, blue light (473 nm) was delivered though the objective using an CoolLED driver (pE-300^ulta^).

Optical stimulation mimicked optical stimulation performed for in vivo electrophysiological recordings. Briefly, 100 stimuli with an ISI of 50 ms (20-Hz) were presented across 10-20 trials with an inter-trial-interval of 25 or 30 seconds. For recordings of optically-evoked inhibitory postsynaptic currents (oIPSCs) cells were voltage clamped at 0 mV (near the theoretical EPSC reversal potential). The maximal peak response and the area under the curve for each stimulus response across all trials was measured and used to generate a stimulus response curve. To assess whether the shape of the stimulus–response curve changed across sweeps, we quantified the similarity between each sweep’s response profile and the first sweep (sweep 0). For each cell, raw AUC (or peak) values across each stimulus were extracted for every sweep. Sweep 0 served as the reference curve. For each subsequent sweep, we computed the Pearson correlation coefficient between that sweep’s stimulus–response vector and that of the first stimulus train. Because correlation is insensitive to absolute amplitude, this metric reflects shape similarity rather than changes in overall response magnitude. Correlation values were computed for every sweep in every cell, then aggregated across cells to obtain the population mean and SEM. To test whether shape similarity changed systematically across sweeps at the population level, we fit a linear mixed-effects model with sweep number as a fixed effect and cell identity as a random effect.

## Results

### Baseline Firing and Subtype Composition of vSub Neurons In Vivo

To begin to assess the role of mDlx⁺ GABAergic interneurons on putative ventral subiculum (vSub) pyramidal neurons, we performed in vivo electrophysiological recordings using multielectrode probes. We performed optical stimulation to phototag GABAergic neurons. Photostimulation occurred over 5 seconds (20-Hz, 1 ms duration) across 20 trials with an intertrial interval of 30 seconds. From a total of 116 well-isolated single units, 16 neurons (∼14%) were classified as GABAergic based on optical tagging (Fig. 1 A-B) and the remaining 100 of these neurons were classified as pyramidal neurons (Fig. 1C).

**Figure 1.**
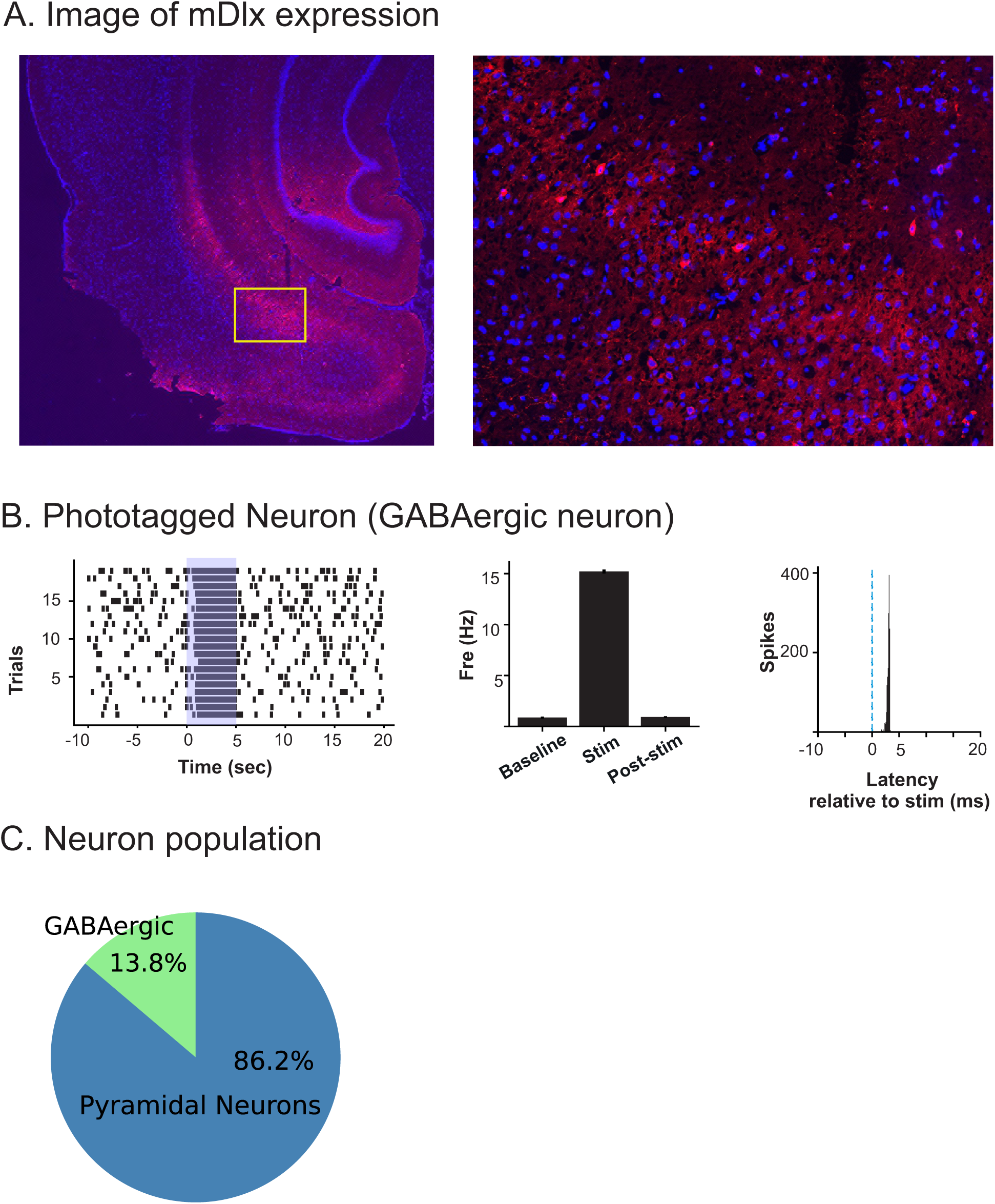
GABAergic neurons in the vSub can be optically identified using the mDlx enhancer element. (A) Representative images showing mDlx⁺ neuron expression in the vHC (left). Yellow box indicates the region shown at higher magnification (right). (B) Spike raster of an optically identified GABA (mDlx⁺) neuron (left), quantification of mean firing rate before, during, and after optical stimulation (center), and peristimulus time histogram aligned (blue dotted line) to stimulation onset (right). (C) Proportion of optically identified GABAergic (mDlx⁺) and putative pyramidal neurons.

### Rebound Excitation Emerges in Pyramidal Neurons Following Optogenetic Inhibition

We next compared the firing properties of the remaining, presumptive, pyramidal neurons. To assess baseline activity we measured the average firing rate during the final ten seconds preceding each optical stimulation of mDlx⁺ neurons. Although we initially assumed that neurons would reliably return to their prestimulus firing rates after each stimulation over the course of the 25 second poststimulus periord, we found a subset of neurons that failed to return to baseline.

To determine whether this deviation reflected a transient post-stimulus rebound or a progressive increase in excitability across repeated stimuli, we quantified spontaneous firing during defined post-stimulation epochs across successive trials. Both cumulative spike counts and trial-averaged firing rates revealed systematic changes in post-stimulus activity over the course of the session. Linear regression of mean post-stimulus firing rate against trial number identified a significant pattern (*p*<0.05) in 38 neurons, indicating a persistent, trial-to-trial change in post-stimulus firing. Of these 38 neurons, 26 showed a significant positive trend (up-rampers) while 12 showed a negative trend (down-rampers) (Fig 2B, C and E).

**Figure 2.**
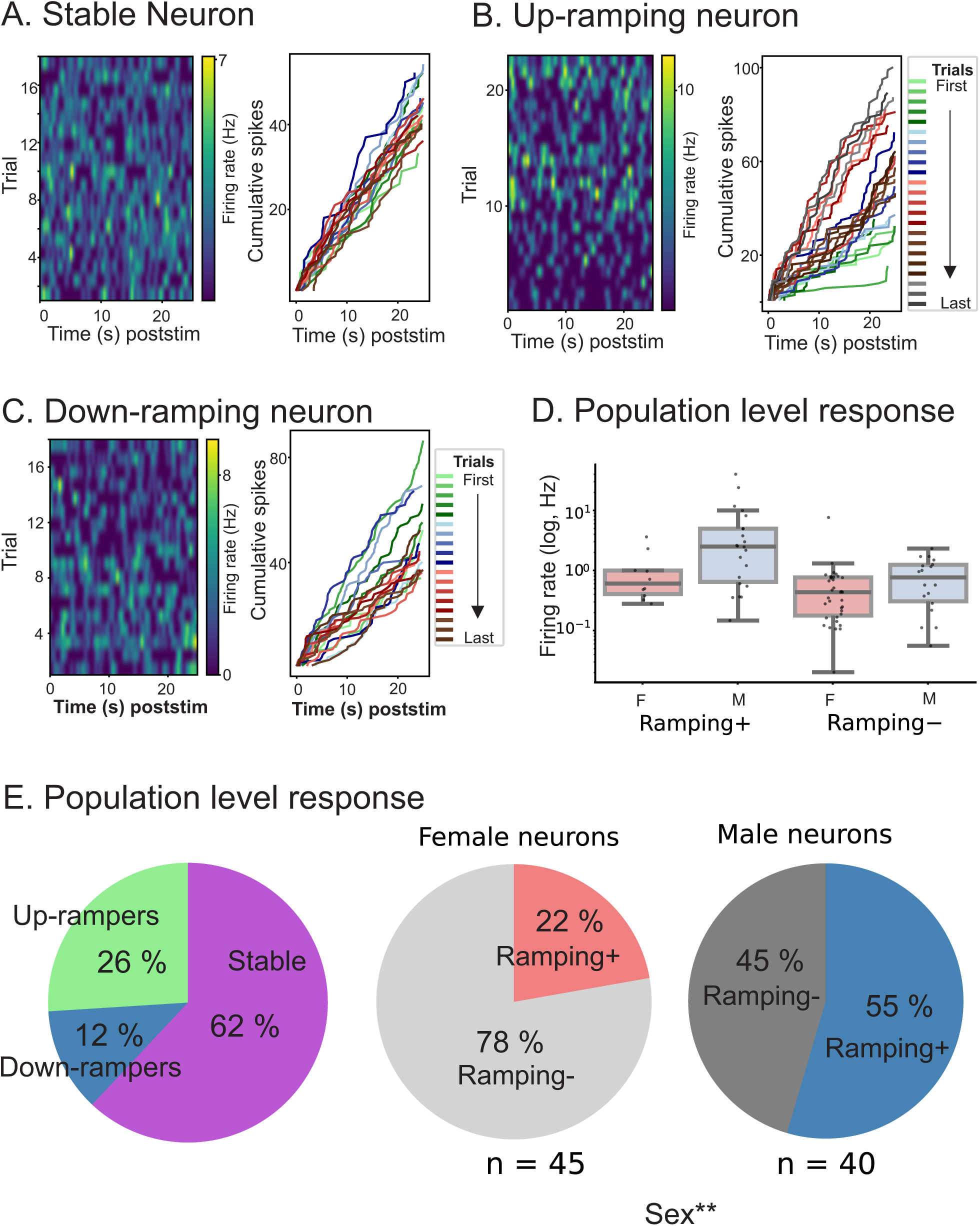
A heterogeneous pattern of poststimulus firing is observed in vSub pyramidal neurons. Representative heat maps (left) and cumulative spike counts (right) are shown for neurons exhibiting (A) stable, (B) up-ramping, or (C) down-ramping poststimulus firing rates. Colors in the cumulative spike count plots represent individual trials, as indicated in the legend. (D) Comparison of firing rates in ramping and non-ramping neurons between male and female rats (D) Proportion of vSub pyramidal neurons exhibiting stable, up-ramping, and down-ramping poststimulus firing patterns and the distribution of ramping and non-ramping neurons in male and female rats. ** indicates p<0.01

These findings suggest that optical inhibition not only produces acute suppression of pyramidal neuron activity but also induces longer-lasting increases in excitability that accumulate across repeated stimulation. When stratified by sex, a significantly greater proportion of male rats exhibited this ramping phenotype compared to their female counterparts (22.2% in females vs 55.0% in males, *χ^2^*(1)= 8.35, *p*=0.004, Fig 2D and E).

Importantly, these divergent response profiles, coexisting within the same recording, argue against a global, nonspecific change in network excitability (for example, a gradual drift in anesthetic depth). The presence of neurons that increased, decreased, or remained stable under identical experimental conditions suggests that post-stimulation changes in spontaneous firing were neuron-specific and likely related to circuit engagement rather than a uniform recording artifact. While this heterogeneity is strongly suggestive of a stimulation-dependent effect, it alone was not sufficient to fully exclude slow, time-dependent confounds.

To more directly address this concern, we analyzed recordings in which a separate baseline recording (5-15 min in duration) was available in the absence of optical stimulation. We performed a permutation analysis in which trial labels were randomly shuffled to generate a null distribution of trial-wise firing rate trajectories. This null distribution was used to determine whether observed firing rate dynamics during stimulation were consistent with systematic monotonic increases or decreases in activity, rather than reflecting random trial-to-trial variability. Using this approach, 56.6% (30 of 53) of neurons showed firing rate trajectories during stimulation that significantly deviated from the null distribution. Among these the majority (80% or 24 of 30) of neurons exhibited ramping increases, while a smaller percentage (20% or 6 of 30) showed a decrease in ramping (Fig 3 A-D).

**Figure 3.**
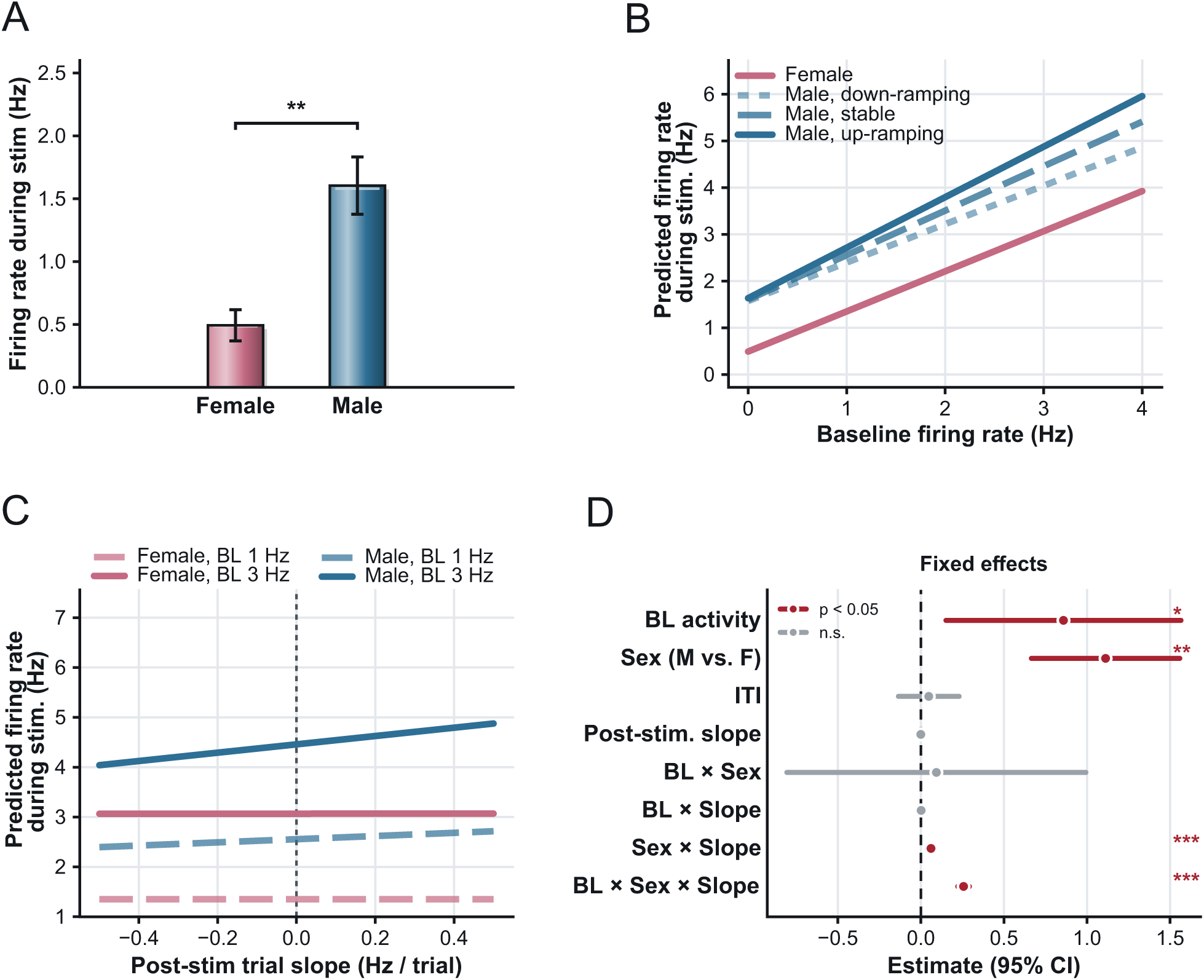
This heterogeneous pattern of poststimulus firing cannot be explained by persistent elevations in firing rates independent of optical stimulation. (A–C, left) Cumulative spike count curves during baseline (gray) and non-stimulated epochs of optically stimulated recordings (red) for neurons exhibiting the three poststimulus firing patterns. (A–C, middle) Permuted null distributions (blue histograms) and observed mean differences in firing rate between baseline and poststimulus epochs (solid red lines). (A–C, right) Cumulative poststimulus spike counts across trials for each neuronal subtype. Colors represent individual trials, as indicated in the legend.

### Effect of Optical Activation of GABAergic Neurons on Networked Pyramidal Neuron Firing

To look more specifically at the direct effect of optical stimulation on pyramidal neuron activity we analyzed pyramidal neuron firing in response to mDlx^+^ neuron-dependent inhibition using a linear mixed-effects model that accounted for the nested dependence structure across trials, neurons, and recordings. Firing rate was treated as the outcome variable and modeled as a function of firing rate period, specified as a categorical predictor with three levels: baseline, stimulation, and post-stimulation.

Because the inter-trial intervals varied across some neurons, a fixed-effects covariate representing trial timing intervals was included to control for its potential confounding effects. To capture within-neuron ramping behavior over successive trials, trial progression (trial number) was included as a continuous fixed effect, along with its interaction with firing rate period, that further allowed examining the influence of ramping on firing rate during each period. Random effects were modeled at two hierarchical levels, including a random intercept at the recording level that captured between-recording variability, and both a random intercept and a random slope (with respect to trial progression) at the neuron level to account for neuron-specific baseline firing rates and unique ramping trajectories across successive trials.

Data from 29 unique recordings and 100 pyramidal neurons were included in this analysis. Results are summarized in Table 1. The estimated effect for stimulation (*Period*^(1)^) was significant (*β* = −1.18, *p*<0.0001), indicating that optical stimulation produced a robust reduction in firing rate across all pyramidal neurons (Fig 4B).

**Figure 4.**
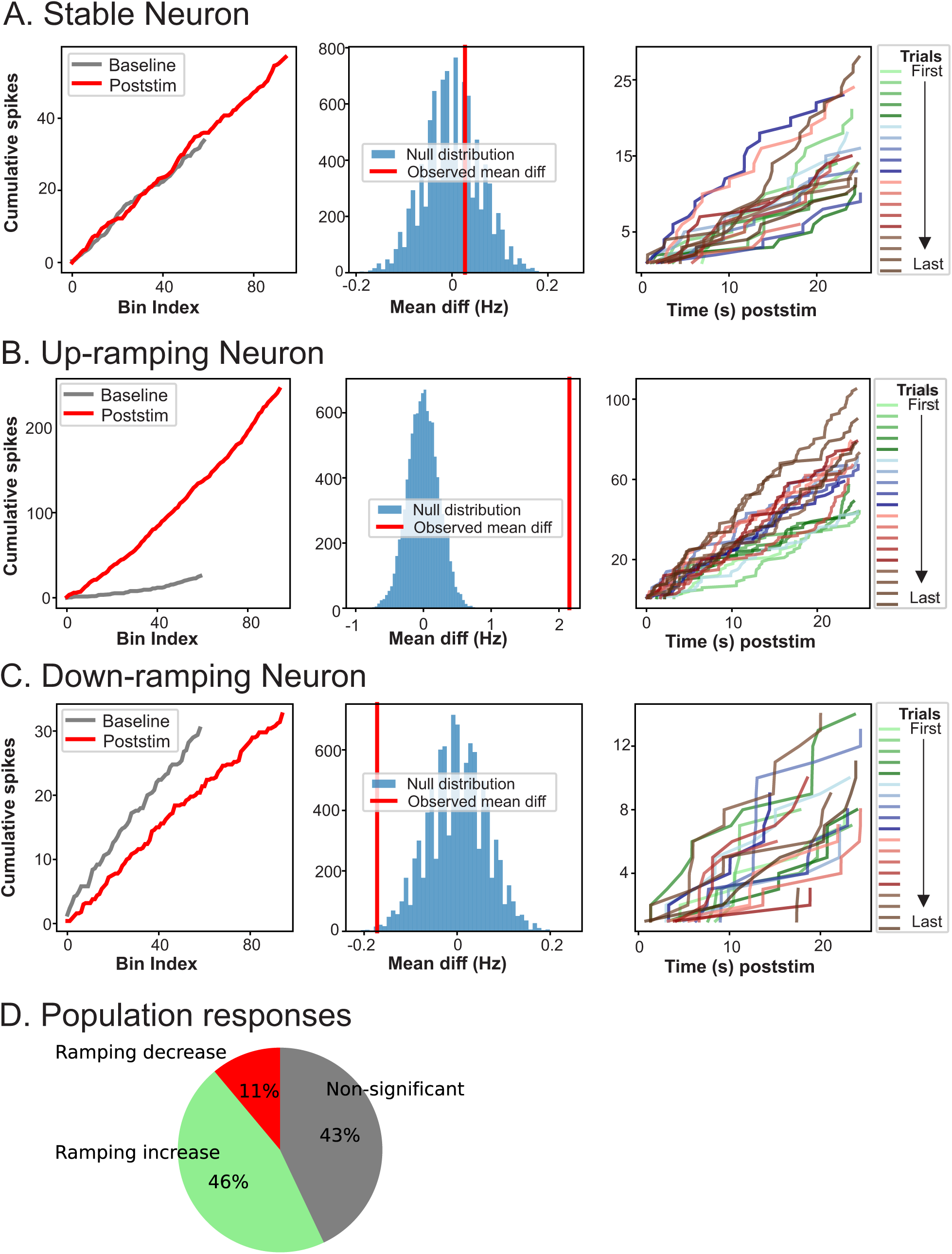
Suppression of pyramidal neuron firing by optical activation of GABAergic neurons. (A) spike raster (left), epoch specific quantification of firing rates (middle) and autocorrelogram (left)of a pyramidal neuron inhibited by optical stimulation. (B) Pyramidal neuron firing rates during baseline, stimulation, and post-stimulation periods. Optical stimulation significantly reduced firing during stimulation, with post-stimulation firing returning to baseline levels. (C) Firing rates during stimulation in early and late trials, showing progressively greater suppression across repeated trials. (D) Firing rates across baseline, stimulation, and post-stimulation periods in male and female rats. (E) Firing rates during early and late stimulation trials in male and female rats, showing greater trial-dependent suppression in males. Values represent estimated marginal means from linear mixed-effects models. Percentages indicate reductions relative to the corresponding baseline. *p < 0.05; **p < 0.01; ***p < 0.001; †p < 0.10; n.s., not significant.

**Table 1.** Firing rate difference between baseline, stimulation, and post-stimulation periods.

|  | **Estimate** | **\*SE** | **\*P-values** |
| --- | --- | --- | --- |
| ***Intercept*** (baseline) | 2.2461 | 0.6412 | 0.0016 |
| ***Period*<sup>(1)</sup>** (stimulation vs.<br>baseline) | -1.1849 | 0.0885 | <0.0001 |
| ***Period*<sup>(2)</sup>** (post-stimulation vs.<br>baseline) | 0.0985 | 0.0891 | 0.3763 |
| ***ITI*** | -0.4358 | 0.4896 | 0.4357 |
| ***Trial*** | 0.4131 | 0.3331 | 0.3761 |
| ***Period*<sup>(1)</sup> × *Trial*** | -0.3521 | 0.1358 | 0.0223 |
| ***Period*<sup>(2)</sup> × *Trial*** | 0.09502 | 0.1380 | 0.4913 |

\*p-values are corrected for multiple comparisons using fdr procedure. \*SE: Standard Error.

Specifically, firing rate decreased by approximately 1.18 Hz (52.7%) relative to baseline, reducing baseline activity from 2.24 ± 0.64 Hz to 1.06 ± 0.64 Hz during stimulation.

The post-stimulation period (*Period*^(2)^) was not significantly different from baseline (*p*=0.3763), with firing rates recovering to nearly the baseline levels (estimated change: 0.10 ± 0.08 Hz). The interaction between stimulation and trial progression (*Period*^(1)^ × *Trial*) was significant as well (*β* = −0.35, *p*=0.0223), indicating that the suppressive effect of optical stimulation increased across successive trials (Fig 4C). According to the estimates in Table 1, firing rates during stimulation decreased by ∼ 0.35 Hz across trials, reaching ∼0.71 Hz in later trials and thus an overall reduction of nearly 68.3% relative to baseline.

To determine whether sex influenced firing rates during optical inhibition and whether it moderated the ramping effects, sex (reference: female) and its interactions with key covariates were included in a subsequent model (Table 2) with the same random effects. After including sex, the reduction in firing rate during optical stimulation was observed in males (p<0.0001) but not in females (*p*=0.31). Among females, the baseline firing rate was estimated at 1.24 ± 0.24 Hz, and decreased by 0.21± 0.13 Hz during stimulation, a change that was not statistically significant. However, males showed a higher baseline firing rate of 3.38 ± 1.28 Hz (derived as 1.24 + 2.14), which decreased by nearly 1.82 ± 0.13 Hz during stimulation (∼ 53.8% reduction)(Fig 4D)

**Table 2.** Influence of Sex on optical stimulation firing rates.

|  | **Estimate** | **\*SE** | **\*P-values** |
| --- | --- | --- | --- |
| ***Intercept*** | 1.2413 | 0.8520 | 0.3164 |
| ***Period*<sup>(1)</sup>** | -0.2046 | 0.1269 | 0.3062 |
| ***Period*<sup>(2)</sup>** | 0.0295 | 0.1278 | 0.8175 |
| ***ITI*** | -0.4307 | 0.4878 | 0.7007 |
| ***Trial*** | -0.1329 | 0.4720 | 0.8176 |
| ***Period*<sup>(1)</sup> × *Trial*** | 0.0663 | 0.1813 | 0.8175 |
| *Period*<sup>(2)</sup> × *Trial* | 0.0462 | 0.1844 | 0.8175 |
| *Sex* | 2.1440 | 1.2774 | 0.3061 |
| *Period*<sup>(1)</sup> × *Sex* | -1.8214 | 0.1741 | <0.001 |
| *Period*<sup>(2)</sup> × *Sex* | 0.1242 | 0.1751 | 0.7771 |
| *Sex* × *Trial* | 1.0376 | 0.6632 | 0.3062 |
| *Period*<sup>(1)</sup> × *Sex* × *Trial* | -0.7043 | 0.2674 | 0.0549 |
| *Period*<sup>(2)</sup> × *Sex* × *Trial* | 0.0926 | 0.2718 | 0.8176 |

\**p*-values are corrected for multiple comparisons using fdr procedure. \*SE: Standard
Error

**Table 3.**
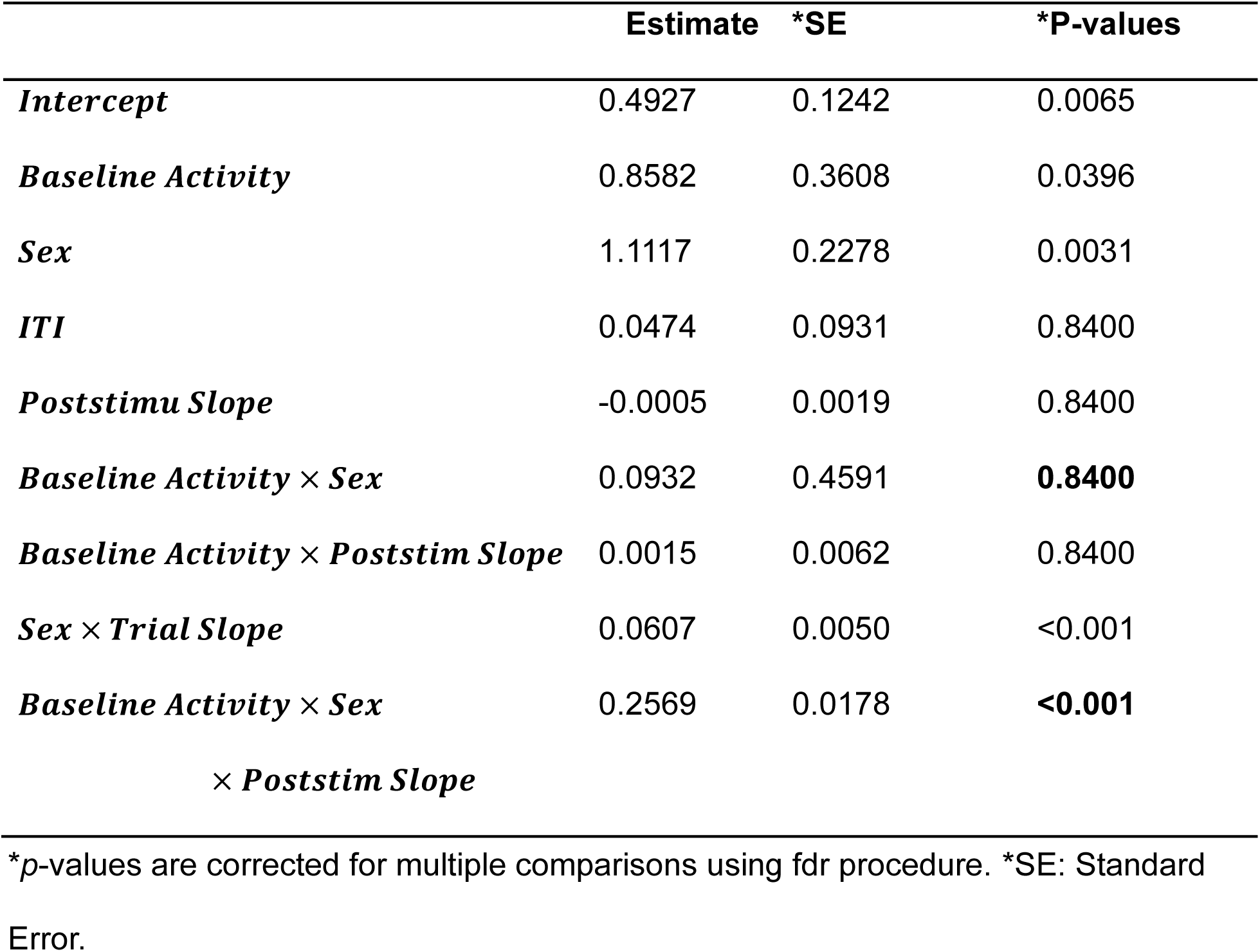
Influence of Sex and ramping behavior on optical stimulation after accounting for baseline activity.

The three-way interaction (*Sex* × *Period*^(1)^ × *Trial*) was also significant, indicating that the suppressive effect of optical stimulation in males increased across successive trials. According to Table 2, firing rates during stimulation decreased by ∼ 0.70 Hz in males during later trials, corresponding to an overall reduction of nearly 74.5% when compared to baseline.

Baseline firing rate was a significant predictor of firing during stimulation (*β* = 0.86 ± 0.36, *p* = 0.0396), indicating that neurons with higher spontaneous activity also maintained higher firing during GABAergic activation. Sex was also a significant predictor, with males showing higher firing rates during stimulation when compared with females (reference group) (*β* = 1.11 ± 0.23, *p* = 0.0031). Specifically, the estimated average firing rate for females during stimulation was 0.49 ± 0.12 *Hz*, and substantially higher rates were observed in males. The interaction between sex and post-stimulus slope (*Sex* × *Trial Slope*) was significant, indicating that greater positive ramping (increasing firing across trials) was associated with higher firing rates during stimulation in males but not females. This effect was further amplified among males with higher baseline firing rates, as reflected in the three-way interaction with baseline activity (Fig 5A-D). Note that the main effect of post-stimulus slope was not significant (*p*=0.8400), likely due to heterogeneity in ramping patterns across neurons, including up-ramping, down-ramping, and stable firing trajectories that may offset one another. These results demonstrate that ramping behavior and sex are important predictors of firing during GABAergic activation and should be considered in interpreting the results.

**Figure 5.**
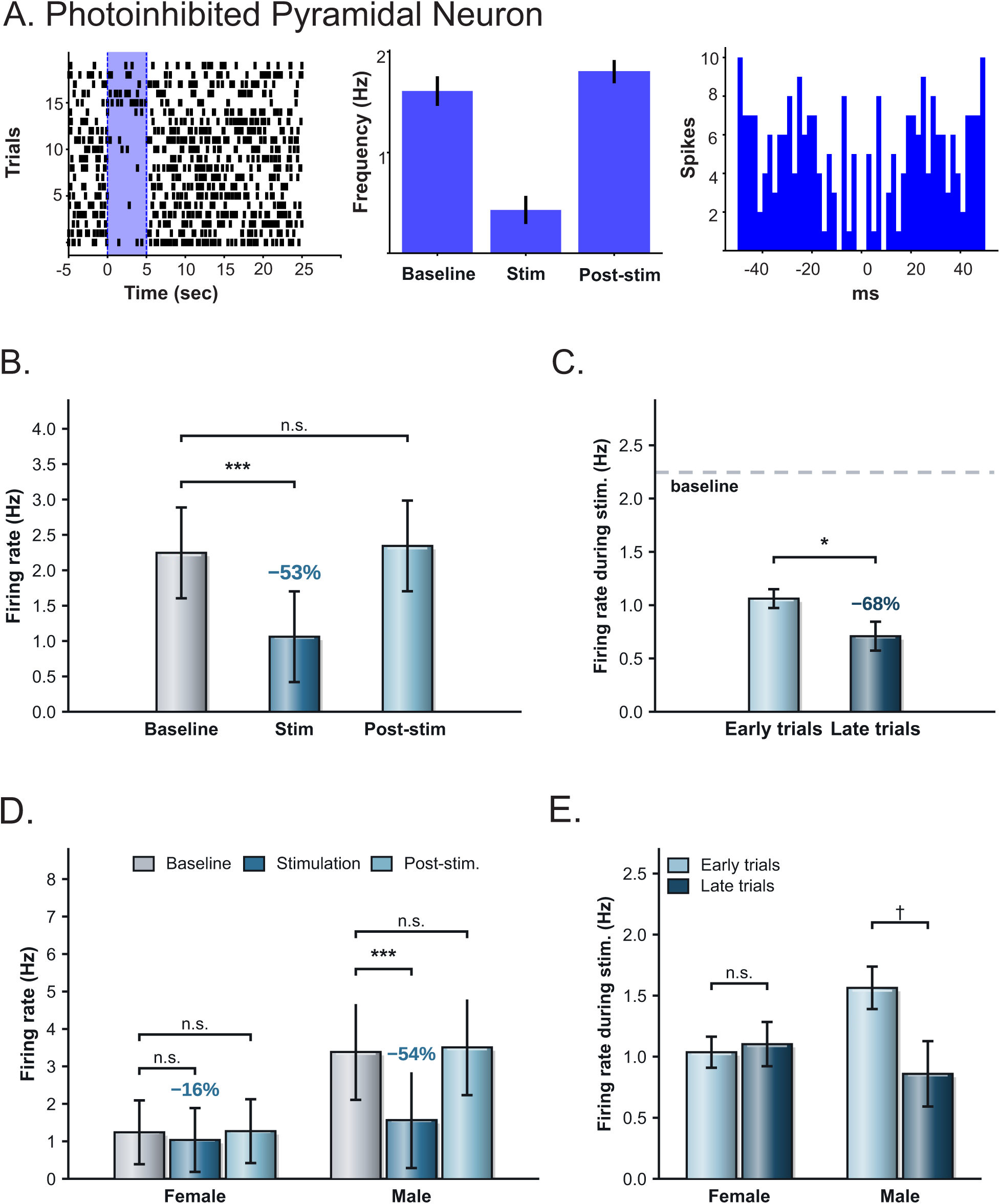
Baseline activity, sex, and ramping behavior predict pyramidal neuron firing during GABAergic activation. (A) Estimated pyramidal neuron firing rates during stimulation in male and female rats. (B) Relationship between baseline firing rate and firing during stimulation, shown across post-stimulation ramping slopes and separated by sex. (C) Relationship between post-stimulation firing slope and firing during stimulation at low and high baseline firing rates in male and female rats. (D) Fixed-effect estimates with 95% confidence intervals from the linear mixed-effects model. The dashed line indicates zero. Values represent model predictions or estimated effects from the linear mixed-effects model. *p < 0.05; **p < 0.01; ***p < 0.001.

Multicollinearity diagnostics showed no problematic collinearity for this analysis (see supplemental file for more detail).

### Neuronal Firing in Relation to Slow-Oscillation Phase

To better assess local network associations, we examined local field potentials (LFPs) and their relationship to neuronal activity during GABAergic stimulation.

Specifically, we assessed the distribution of action potentials across the phase of the slow oscillation before, during, and after stimulation to determine whether phase shifts might contribute to population-dependent differences in ramping behavior. A striking feature that emerged was that a majority of GABAergic neurons exhibited evidence of phase preference to the slow oscillation during the prestimulus epoch (10/16, 63%), whereas a smaller proportion of pyramidal neurons showed similar phase preferences (41/100, 41%). To further assess changes in phase preferences across stimulation epochs, we restricted this analysis to neurons with a minimum of 10 action potentials in each epoch (before, during, and after stimulation). Among GABAergic neurons, stimulation was associated with a pronounced change in phase preference, with nearly all significant changes reflecting a loss of phase locking during stimulation relative to the prestimulus epoch (8/12, 66.7%, Fig 6A-B) and after stimulation (9/12, 75.0%). In contrast, substantially fewer pyramidal neurons exhibited significant changes in phase preference during stimulation relative to either the prestimulation epoch (7/56, 12.5% Fig 6A-B) or the post-stimulation epoch (9/56, 16.1%). To begin to assess whether the ramping behavior of pyramidal neurons may be related to changes in phase locking, we examined the relationship between ramping and both the strength of phase locking and changes in phase locking across stimulation epochs. None of these measures were significantly correlated with one another (all Spearman correlations, p > 0.05), suggesting that neither the strength of phase locking nor changes in phase locking are prominent contributors to the observed ramping behavior.

**Figure 6.**
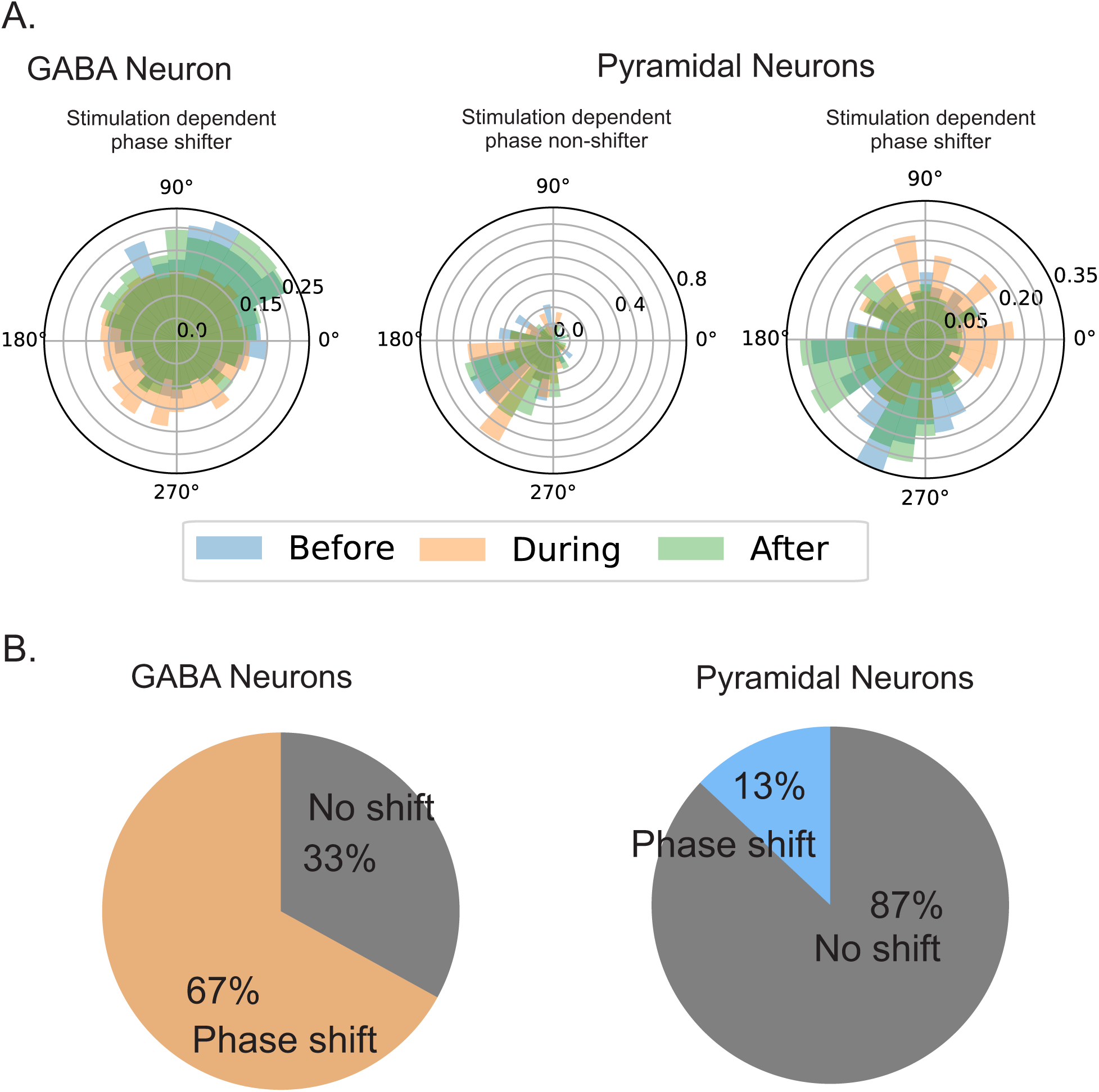
A greater proportion of GABAergic neurons exhibited stimulation-dependent alterations in phase preference compared with pyramidal neurons. (A) Representative example of a phase-preferring GABA neuron showing a loss of phase preference during stimulation. (B) Representative phase-preferring pyramidal neurons showing no change (left) or a shift (right) in phase preference in response to local GABA neuron stimulation. (C) Population-level distributions of stimulation dependent phase shift behavior in GABA (left) and pyramidal (right) neurons.

### Progressive Changes in Magnitude and Temporal Structure of Inhibitory Post Synaptic Currents

To begin to investigate synaptic mechanisms that may contribute to changes in rebound excitation observed in vivo, we performed ex vivo whole-cell recordings from pyramidal neurons in the ventral subiculum of animals expressing channelrhodopsin (ChR2) in mDlx^+^ GABAergic neurons. This approach allowed direct measurement of optically evoked inhibitory postsynaptic currents (oIPSCs) during repeated stimulation paradigms identical to those used for the in vivo electrophysiology experiments. Across neurons, oIPSC responses declined systematically over successive trials. Mixed-effects modeling revealed a highly significant effect of sweep number on response magnitude (β = −1.54 ± 0.22 per sweep, p < 0.0001; Fig 7), indicating a progressive reduction in synaptic inhibitory strength (Fig 7 A-C, F).

**Figure 7.**
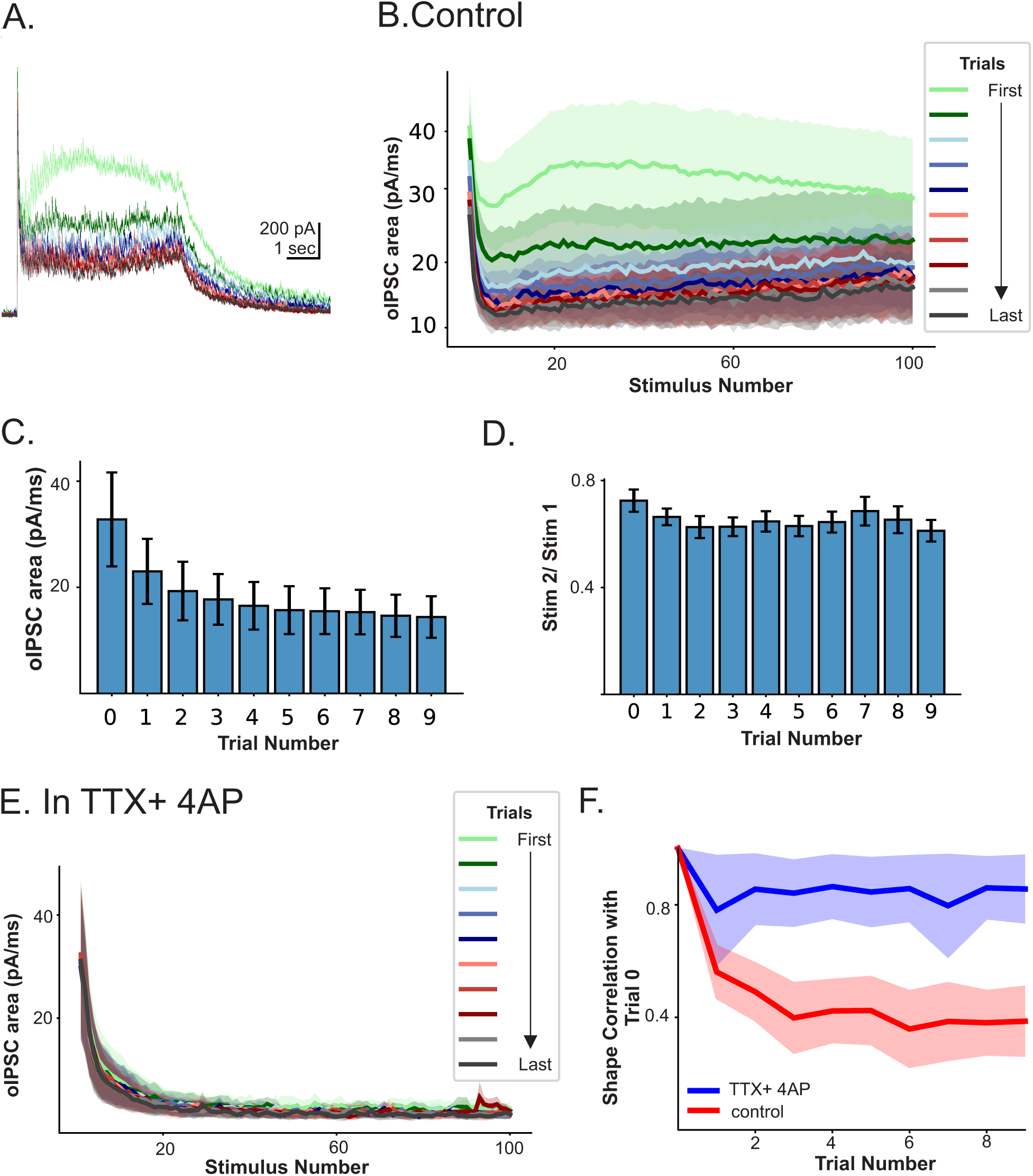
Repeated trials of 20 Hz stimulation elicit short-term plasticity that requires network engagement. (A) Representative oIPSCs in a vSub pyramidal neuron presented with GABA neuron optical stimulation (20 Hz, 100 stimuli) over the course of 10 stimuli. (B) Group oIPSC response to optical stimulation across trials under control conditions (rACSF). In A and B colors represent individual trials, as indicated in the legend. In B areas shaded in light trial colors indicate SEM of group response. (C) Quantification of group oIPSC areas in pyramidal neurons during each optically stimulated trial. (D) Quantification of group comparisons of the first and second stimulus (PPr) within and across trials. Error bar represent ±SEM. (E) Group oIPSC response to optical stimulation across trials under in the presence of TTX+4AP. (F) Comparison of the trial dependent shape correlation between neurons recorded in rACSF and TTX+4AP.

Changes in synaptic responses were not limited to overall magnitude. A mixed-effects model revealed a significant negative relationship between sweep number and paired-pulse ratio (slope = −0.007 ± 0.002 per sweep, *p*=0.002; intercept = 0.68 ± 0.03, Fig 7 D), consistent with altered short-term synaptic dynamics across repeated stimulation. Because paired-pulse ratio is sensitive to presynaptic release mechanisms, this result suggests that repeated stimulation modifies release probability or vesicle availability rather than reflecting uniform rundown of synaptic strength. To examine whether repeated stimulation also altered the temporal structure of inhibitory responses, we performed shape-correlation analyses comparing the pattern of oIPSCs within a trial across successive sweeps. Shape correlations relative to the first sweep declined progressively across trials. A mixed-effects model revealed a significant negative relationship between sweep number and shape correlation (slope = −0.049 ± 0.007 correlation units per sweep, *z* = −6.93, *p* < 0.001; intercept = 0.689 ± 0.102). These results indicate that repeated stimulation not only reduces the magnitude of inhibition, but also systematically alters the relative timing and structure of inhibitory input independent of absolute amplitude (Fig. 7 A-C, F).

To test whether engagement of circuit-level interactions contributes to progressive response degradation, we applied TTX and 4-AP to restrict activity to presynaptic terminal release. In the presence of TTX+4AP there was no progressive change in response shape across stimulus trains (β = −0.006, *p*=0.297, Fig 7 E, F). Mixed-effects modeling revealed a significant interaction between sweep number and condition (β = −0.040 ± 0.017, *p*=0.020), indicating that response shape degraded progressively across stimulus trains in control neurons but remained stable in the presence of TTX+4AP (Fig 7).

## Discussion

Our findings demonstrate that optogenetic activation of mDlx⁺ GABAergic interneurons within the vSub elicits robust inhibition of pyramidal neurons during optical stimulation, confirming effective recruitment of local inhibitory circuitry. To the best of our knowledge, the current study represents the first examination of the functional consequences of recruiting the broader GABAergic neuronal population within the vSub, extending previous studies that have focused on more restricted interneuron populations (Bratsch-Prince et al., 2026; Forro et al., 2022; Volitaki et al., 2024). Apart from this direct effect, however, we further establish that the activation of mDlx⁺ GABAergic neurons engages circuit mechanisms that extend well beyond the period of direct inhibition. Repeated stimulation trials revealed that the inhibitory response was not physiologically static. Although many pyramidal neurons returned to their baseline firing rates following each stimulation epoch, representing stable poststimulus activity across trials, a substantial proportion of neurons exhibited poststimulus dependent firing rate adaptations. Of these adapting neurons, the majority increased their firing rate (up-rampers) while a minority decreased their firing rate (down-rampers) across trials. At the same time, the magnitude of inhibition during optical stimulation itself progressively increased over repeated trials but this finding was heavily influenced by both sex and poststimulus firing rates. Together, these findings indicate that repeated activation of vSub mDlx⁺ interneurons simultaneously strengthens acute inhibition while producing longer-lasting changes in spontaneous network excitability, suggesting that distinct mechanisms govern activity during and after optical stimulation.

The observation that repeated stimulation altered neuronal activity well beyond the period of optical inhibition suggests that recruitment of inhibitory circuits initiates secondary network processes that evolve over successive stimulation epochs.

Importantly, these effects were not uniform across the population, as neurons displaying increasing, decreasing, or stable firing trajectories were often observed within the same recording. This heterogeneity, together with permutation analyses using extended baseline recordings, indicates that the observed ramping behavior is unlikely to reflect recording instability or gradual changes in anesthetic depth, but instead represents a stimulation-dependent recruitment of network activity. Consistent with this finding, baseline firing rate predicted the magnitude of inhibition in individual pyramidal neurons. Because baseline firing rate reflects the ongoing physiological state of the recorded neuron within the local network, rather than the level of ChR2 expression in mDlx⁺ neurons, this relationship suggests that variability in inhibitory efficacy is more likely to arise from differences in the ongoing state of circuit engagement rather than from differences in opsin expression or optical recruitment. Thus, the effectiveness of GABAergic recruitment appears to be influenced by the pre-existing network state of the recorded neuron. The extensive connectivity among and between neuronal subtypes within the vSub, including pyramidal and GABAergic neurons, provides a network architecture through which recruitment of mDlx⁺ interneurons could engage secondary network interactions and produce heterogeneous changes in pyramidal neuron activity over successive stimulation epochs (Böhm et al., 2015; K. Li et al., 2024; Wee and MacAskill, 2020).

The role of sex in our findings is an interesting one, but one that must be understood with some level of ambiguity. Although sex was a significant predictor of firing during stimulation, females in our dataset exhibited substantially lower baseline firing rates, and baseline firing rate was itself a strong predictor of firing during stimulation. Moreover, the interaction between baseline activity and ramping indicates that the response to repeated GABAergic activation is influenced by the pre-existing activity state of the neuron. Thus, the observed difference between males and females cannot be readily interpreted as evidence for an intrinsic sex-dependent difference in the response to GABAergic recruitment. Rather, the lower baseline activity observed in females may contribute to, or constrain, their firing response during stimulation and the extent to which activity changes across successive trials. Consistent with this possibility, ramping responses were disproportionately represented among males, suggesting that sex-associated differences may extend beyond the immediate response to stimulation to influence how neuronal activity evolves across repeated recruitment. Importantly, because baseline activity, sex, and ramping behavior are not independent features of these data, the present experiments cannot determine whether males and females with comparable baseline firing rates would exhibit equivalent responses to repeated GABAergic activation. To the best of our knowledge, sex differences in the basal firing rates of pyramidal neurons have not been reported. However, this is difficult to assess definitively, as many in vivo studies have been conducted exclusively in males (Forro et al., 2022; Forro and Klausberger, 2023; Simonnet and Brecht, 2019), while others do not report whether statistical comparisons of basal firing rates between sexes were performed (K. Li et al., 2024). Given that our studies were conducted under anesthetic conditions, differential sensitivity to the anesthetic between males and females also cannot be ruled out as a contributing factor to the observed difference in baseline firing. Despite this ambiguity we consider this to be an important relationship between pre-existing neuronal activity and the subsequent response to repeated GABAergic recruitment.

Given our proposed engagement of local network circuitry, we also sought to determine whether GABAergic recruitment altered the organization of ongoing network oscillations. Because our recordings were performed under sodium pentobarbital anesthesia, our analysis of oscillatory activity was limited to slow oscillations, which are retained across a variety of anesthetic states, including sodium pentobarbital anesthesia (Steriade et al., 1993), and have been implicated in cortical communication and long-range network coordination, in part through their interactions with other oscillatory rhythms (Espinosa-Martínez et al., 2025; Mattia and Sanchez-Vives, 2012; Staresina, 2024). Given the disproportionate number of GABAergic neurons exhibiting evidence of phase locking and associated loss of phase locking during GABAergic stimulation (perhaps not unexpectedly given the direct optical activation at phase-independent onsets), we sought to determine whether disruption of local phase structure might contribute to subsequent changes in neuronal activity, including the observed ramping response of pyramidal neurons. However, we found no evidence for such a relationship, as the extent of poststimulus ramping was not correlated with measures of phase change. It remains possible, however, that alterations in phase organization influence subsequent firing specifically within optically recruited GABAergic neurons and their pyramidal neuron partners, a relationship that may be obscured when phase and firing responses are examined at the population level rather than within established neuronal pairs. We intend to explore this possibility in future studies using high-density multielectrode probes, which may provide greater opportunity to resolve these relationships at the level of interconnected neurons.

The *ex vivo* electrophysiological experiments provide important insight into the potential mechanisms underlying the progressive network adaptations observed in vivo by enabling direct assessment of how repeated mDlx⁺ interneuron activation influences synaptic inhibition onto hippocampal pyramidal neurons and overall support the significance of network interactions. Repeated optical stimulation of mDlx⁺ interneurons resulted in a progressive reduction in the magnitude of oIPSCs recorded from pyramidal neurons. While this reduction is consistent with a potential contribution from GABA_A_ receptor desensitization (Bianchi et al., 2001; Chang et al., 2002), a postsynaptic mechanism, this explanation alone does not fully account for the additional changes observed following repeated stimulation. Consistent with presynaptic contributions, repeated activation produced changes in paired-pulse ratio, suggesting alterations in release probability or other aspects of presynaptic dynamics during repeated trial activation (Dobrunz and Stevens, 1997; Zucker and Regehr, 2002). Additionally, repeated stimulation produced a trial-dependent alteration in the temporal profile of the inhibitory response, with differences observed between the initial stimulus and subsequent trials, revealing changes beyond a reduction in response magnitude. These alterations in response shape were observed only when the local circuit remained intact and were abolished by TTX and 4-AP, demonstrating that they depend on action potential-dependent network activity rather than intrinsic properties of the directly stimulated synapses. Thus, the progressive reduction in IPSC amplitude and alterations in inhibitory response dynamics likely reflect a complex interplay between presynaptic release mechanisms, postsynaptic receptor responsiveness, and network-dependent circuit interactions rather than a single process underlying synaptic adaptation. While these studies provide some initial insight into this complex interplay, additional studies will be needed to delineate underlying molecular and circuit mechanisms to account for these changes.

Optical stimulation at 20-Hz is widely used in behavioral studies to activate both pyramidal (Felix-Ortiz et al., 2013; Felix-Ortiz and Tye, 2014) and GABAergic neurons (M. Li et al., 2024; Ohta et al., 2022; Tian et al., 2018), Our *in vivo* and *ex vivo* findings add to a growing body of literature indicating that the physiological and behavioral consequences of temporal stimulus dynamics, influenced by both the frequency of stimulation as well as the kinetic properties of the specific opsin should be considered in the context of its broader network effects (Baleisyte et al., 2022; Ohta et al., 2022; Pennington et al., 2024; Strüber et al., 2022; Volitaki et al., 2024). Our findings demonstrate that repeated 20-Hz recruitment of GABAergic neurons can produce heterogeneous changes across pyramidal neuron populations that extend beyond the stimulation period, including persistent changes in firing and the emergence of ramping responses across successive stimulation epochs. These prolonged and heterogeneous responses may, in part, reflect the engagement of multiple short term plasticity mechanisms as suggested by our *ex vivo* findings. These findings provide mechanistic links between repeated GABAergic recruitment and the evolving network responses observed *in vivo*. Such complex and heterogeneous responses may have important implications for behavioral studies, where the resulting behavioral phenotype may reflect not only the immediate effects of neuronal activation but also secondary network interactions and longer-lasting changes in circuit activity. This may be particularly relevant when interpreting diverse behavioral outcomes, as different behaviors may engage distinct circuit populations and therefore be differentially sensitive to these heterogeneous and temporally extended effects of stimulation.

Beyond AUD specifically, these findings point to a broader principle relevant to vSub circuit-level dysfunction noted in the neurological/neuropsychiatric disorders referenced in the Introduction: repeated recruitment of the vSub’s mDlx^+^ GABAergic population produces a circuit state that evolves rather than remains fixed. As one example, the subiculum is a well-established site of seizure initiation in temporal lobe epilepsy, and loss of GABAergic inhibition has been directly linked to the emergence of epileptiform activity (Knopp et al., 2008). Our finding that this population’s acute inhibitory efficacy strengthens even as it produces rebound hyperexcitability that outlasts stimulation offers a candidate mechanism by which an intact inhibitory circuit could nonetheless destabilize under repetitive engagement. Because the magnitude of this response was determined primarily by each neuron’s pre-existing activity state rather than by sex, these data further suggest that circuit-based interventions may need to account for baseline network state, a consideration increasingly emphasized in the design of state-adaptive and closed-loop neuromodulation approaches (Sellers et al., 2024). Within the specific context of AUD, the long-term adaptations in the vSub that result as a consequence of ethanol exposure (Bach et al., 2021a, 2021b; Bach and Weiner, 2025; Ewin et al., 2019) may be particularly consequential given that the hippocampus not only contributes to the representation of contextual and affective information, but also retains the capacity to flexibly update the value assigned to stimuli (Biane et al., 2025). Thus, ethanol-induced changes within a circuit capable of dynamically updating stimulus value may influence how ethanol-associated and anxiety-provoking experiences are subsequently represented and assigned motivational significance. In light of our identification of robustly strengthened GABAergic neurotransmission, which may represent an early component of binge-drinking-associated plasticity (Bach et al., 2021b, 2021a; Bach and Weiner, 2025; Ewin et al., 2019), an important speculative question is whether repeated GABAergic recruitment produces alterations in network activity that could contribute to, or set the stage for, longer-term plasticity within the ventral hippocampus associated with AUD. Future studies will be aimed at exploring this possibility.

Taken together, these findings suggest that optogenetic activation of mDlx⁺ interneurons does more than transiently suppress pyramidal neuron firing. Instead, repeated stimulation appears to recruit dynamic circuit processes that reshape both neuronal excitability and the temporal organization of network activity. These network-level adaptations likely include disinhibitory and recurrent circuit interactions that evolve across repeated stimulation and persist beyond the period of light delivery.

Consequently, stimulation paradigms commonly used in behavioral optogenetic studies may not produce identical physiological effects throughout an experiment but instead induce cumulative changes in circuit function that should be considered when interpreting behavioral outcomes.

